# GABA_A_ receptor-mediated currents and hormone mRNAs in cells expressing more than one hormone transcript in intact human pancreatic islets

**DOI:** 10.1101/635144

**Authors:** Sergiy V. Korol, Zhe Jin, Bryndis Birnir

**Author notes:** **Corresponding author**: Dr. Sergiy V. Korol.

## Abstract

In pancreatic islets the major cell-types are α, β and δ cells, secreting the hormones glucagon (GCG), insulin (INS) and somatostatin (SST), respectively. The GABA (γ-aminobutyric acid) signalling system is expressed in human pancreatic islets. We have previously used single-cell RT-PCR in combination with current recordings to correlate expression of single hormone transcript with functional <u>GABA</u>_<u>A</u>_ receptor (<u>iGABA</u>_<u>A</u>_<u>R</u>) properties in islets. Here we extended these studies to islet cells from non-diabetic and type 2 diabetic donors that express mRNAs for more than one hormone. We detected cells expressing double (α/β, α/δ, β/δ cell-types) and triple (α/β/δ cell-type) hormone transcripts. The most common mixed-identity cell-type was the α/β group where the cells could be grouped into β- and α-like subgroups. The β-like cells had low *GCG/INS* expression ratio (< 0.6) and significantly higher frequency of single-channel iGABA_A_R openings than the α-like cells where the *GCG/INS* expression ratio was high (> 1.2). The difference in expression levels and single channel iGABA_A_R characteristics varied in the α/β/δ cell-type. No correlation was observed between the cell-types identity with time in culture or cell size. Clearly, multiple hormone transcripts can be expressed in islet cells whereas iGABA_A_R functional properties appear α or β cell specific.

## Introduction

The three major cell types of the endocrine pancreas are α, β and δ cells [1], producing glucagon, insulin and somatostatin, respectively. When the physiological or pathological aspects of pancreatic islets are studied, the function of α or β cells is traditionally in the focus. However, emerging evidence indicates there are subgroups of pancreatic islet cells that previously were overlooked [2, 3]. Among these are groups of cells expressing more than one hormone transcript [4-6]. They may express hormone transcripts in different combinations such as *GCG/INS, INS/SST, GCG/SST* or *GCG/INS/SST* and have different levels in individual cells. Such cells are here termed “mixed-identity cells”. These cells may potentially represent different developmental stages of the primary cell types [1, 7] but also may appear as a consequence of exposure to different conditions, e.g. development of obesity or diabetes [4, 5].

Elements of the different neurotransmitter signalling machineries are found within human pancreatic islets and one of them is the GABA signalling system [8-11]. This system has been shown to modulate exocytosis [10], insulin and glucagon secretion [8, 9] and regulate β cell replication [11, 12]. In addition, the GABA_A_ receptors in β cells in intact human pancreatic islets and their functional properties have been recently characterized in details [10]. Here we examined the prominence of the single and multiple hormone transcript-expressing cells within intact human pancreatic islets from non-diabetic and type 2 diabetic donors, examined patterns of activity of iGABA_A_Rs in the mixed-identity cells and correlated the channel characteristics with the hormones’ mRNA ratios. Together, the results identify the iGABA_A_Rs as a functional marker of the physiological identity of the mixed-identity cell subtype.

## Methods

### Intact human islets of Langerhans

The Nordic Network for Clinical Islet Transplantation generously provided human pancreatic islets. All procedures were approved by the regional ethics committee in Uppsala (Sweden). Experiments were carried out in accordance with the guidelines and regulations stipulated by appropriate Swedish and European legislation and informed consent was obtained from donors or their relatives. The pancreata from non-diabetic and type 2 diabetic donors were treated by collagenase, and the islets were isolated by Biocoll gradient centrifugation [13]. After that the islets were picked and cultured in CMRL 1066 (ICN Biomedicals, Costa Mesa, CA, USA) with the addition of 10 mM HEPES, 2 mM L-glutamine, 50 μg/ml gentamicin, 0.25 μg/ml fungizone (GIBCO, BRL, Gaithersburg, MD, USA), 20 μg/ml ciprofloxacin (Bayer Healthcare, Leverkusen, Germany), and 10 mM nicotinamide at 37 °C in a high-humidity atmosphere containing 5 % CO_2_, vol/vol and used in the experiments from the second up to fourteen day of culturing.

### Electrophysiological recordings

The electrophysiological recordings from cells in the superficial layers in intact islets were done in the whole-cell patch-clamp configuration using the blind approach. The intact islet was held by the wide-bore holding pipette, and the cell within the islet was approached by the recording pipette from the opposite side. The composition of extracellular solution (in mM): 137 NaCl, 5.6 KCl, 2.6 CaCl_2_, 1.2 MgCl_2_, 10 HEPES and 20 glucose (pH 7.4 using NaOH). The high glucose concentration enhances the vesicular release [14], and we used this phenomenon to stimulate GABA release from the β cells and thus maximize the interstitial GABA concentration within the islets in our experiments in order to facilitate the detection of the GABA_A_ receptor activity. The intracellular solution consisted of (mM): 135 CsCl, 30 CsOH, 1 MgCl_2_, 10 EGTA, 5 HEPES and 3 Mg-ATP (pH 7.2 with HCl). Drugs were purchased from Sigma-Aldrich (Steinheim, Germany) or Ascent Scientific (Bristol, UK). Recordings were done using an Axopatch 200B amplifier, filtered at 2 kHz, digitized on-line at 10 kHz using an analog-to-digital converter and Clampex 10.5 (Molecular Devices, San Jose, CA, USA) software. The access resistance was monitored and if it changed by more than 25%, the recording was rejected.

### Cytoplasm harvesting and single-cell RT-PCR

The cytosome harvesting procedure and single-cell RT-PCR was previously described [10, 15]. Briefly, after completing the patch-clamp experiment in the whole-cell configuration, the negative pressure was applied to the back of the pipette and was relieved at the moment of the whole-cell configuration destroying, and then the pipette content was locked at the atmospheric pressure. These manipulations allowed to collect the cytosome from the cell the electrophysiological recording was done from. The pipette content (5 μL) was expelled to a 200-μL RNase-free PCR tube. The collected cytosome was subjected to the reverse transcription (RT) performed with Verso™ cDNA synthesis kit (Thermo Scientific Waltham, MA, USA). The 20 μL of RT-reaction was exposed to 42 °C for 30 min and then incubated at 95 °C for 2 min. PCR was accomplished according to a standard procedure [10]. The primers for hormone transcripts are glucagon (forward: GCAACGTTCCCTTCAAGACAC, reverse: ACTGGTGAATGTGCCCTGTG), insulin (forward: CCATCAAGCAGATCACTG, reverse: CACTAGGTAGAGAGCTTCC), and somatostatin (forward: CCCAGACTCCGTCAGTTTCT, reverse: AAGTACTTGGCCAGTTCCTGC). The efficiency of primers for each hormone transcript was in the range between 99 and 100%. The relative expression of pairs of hormone transcripts (mRNA) in individual mixed-identity cells was defined as 2^-(Ct[mRNA1]-Ct[mRNA2])^. The melting curve of the PCR product was examined and/or PCR product was run on a 1.5% agarose gel. RNA from whole human islet samples and the intracellular solution or water served as the positive control and negative control, respectively.

### Data analysis

Statistical dependences between different parameters measured in electrophysiological or single-cell RT-PCR experiments were tested by Spearman correlation using GraphPad Prism 7 (La Jolla, CA, USA). The Tukey method was used for the detection of outliers which were excluded from the analysis. Nonparametric Mann-Whitney test was used to compare groups which contained not normally distributed data. Significance level was set at P < 0.05. The values are mean ± S. E. M.

## Results

### Cell-types identified by hormone mRNA expression in intact pancreatic islets from non-diabetic and type 2 diabetic donors

GABA-activated single-channel currents were detected in 383 cells in intact islets from 109 donors. The cell-type was determined by single-cell RT-PCR analysis of the levels of islet insulin (*INS*), glucagon (*GCG*) and somatostatin (*SST*) transcripts for every individual cell recorded from. Hormone transcripts were detected in 174 cells from 45 non-diabetic and 8 type 2 diabetic donors (HbA1c = 6.5 ± 0.16, mean ± S.E.M. (48 mmol/mol)). Table 1 shows the distribution of the cell-types identified. Characteristics of GABA-activated currents in the α, β and δ single-hormone cell-types have been described [10]. Here we analysed the samples containing multiple hormone transcript-expressing cells. For islets from non-diabetic and type 2 diabetic donors, single-hormone transcript was detected in 55% and 48% of the cells, respectively, with 44% (non-diabetic) and 32% (type 2 diabetic donors) of the cells being insulin-positive β cells (Fig. 1A). The remaining cells, 45% from non-diabetic and 52% from type 2 diabetic donors, were positive for more than one hormone transcript. The frequency of the specific subtypes of mixed-identity cells i.e. α/β, β/δ, α/δ, α/β/δ, varied somewhat between the non-diabetic and type 2 diabetic donor islets, with the most notable difference being a decrease in β/δ and an increase in mixed-identity cell subtypes expressing the *GCG* in type 2 diabetic donors (Fig. 1A, Table 1). As the data from type 2 diabetic donors was limited and overlapped with the data from the non-diabetic donors, we combined the results from the two groups when examining single-channel properties and effects of days in culture on the channel properties (Fig. 2, 3).

**Table 1.** Cell-types identified based on expression of hormone mRNAs in pancreatic islets from non-diabetic and type 2 diabetic donors.

| Cell type | $\alpha$ | $\beta$ | $\delta$ | $\alpha/\beta$ | $\beta/\delta$ | $\alpha/\delta$ | $\alpha/\beta/\delta$ | Total |
| --- | --- | --- | --- | --- | --- | --- | --- | --- |
| **Non-diabetic islets** |  |  |  |  |  |  |  |  |
| n cells | 12 | 65 | 4 | 34 | 18 | 2 | 14 | 149 |
| **Type 2 diabetic islets** |  |  |  |  |  |  |  |  |
| n cells | 3 | 8 | 1 | 8 | 1 | 1 | 3 | 25 |

**Figure 1.**
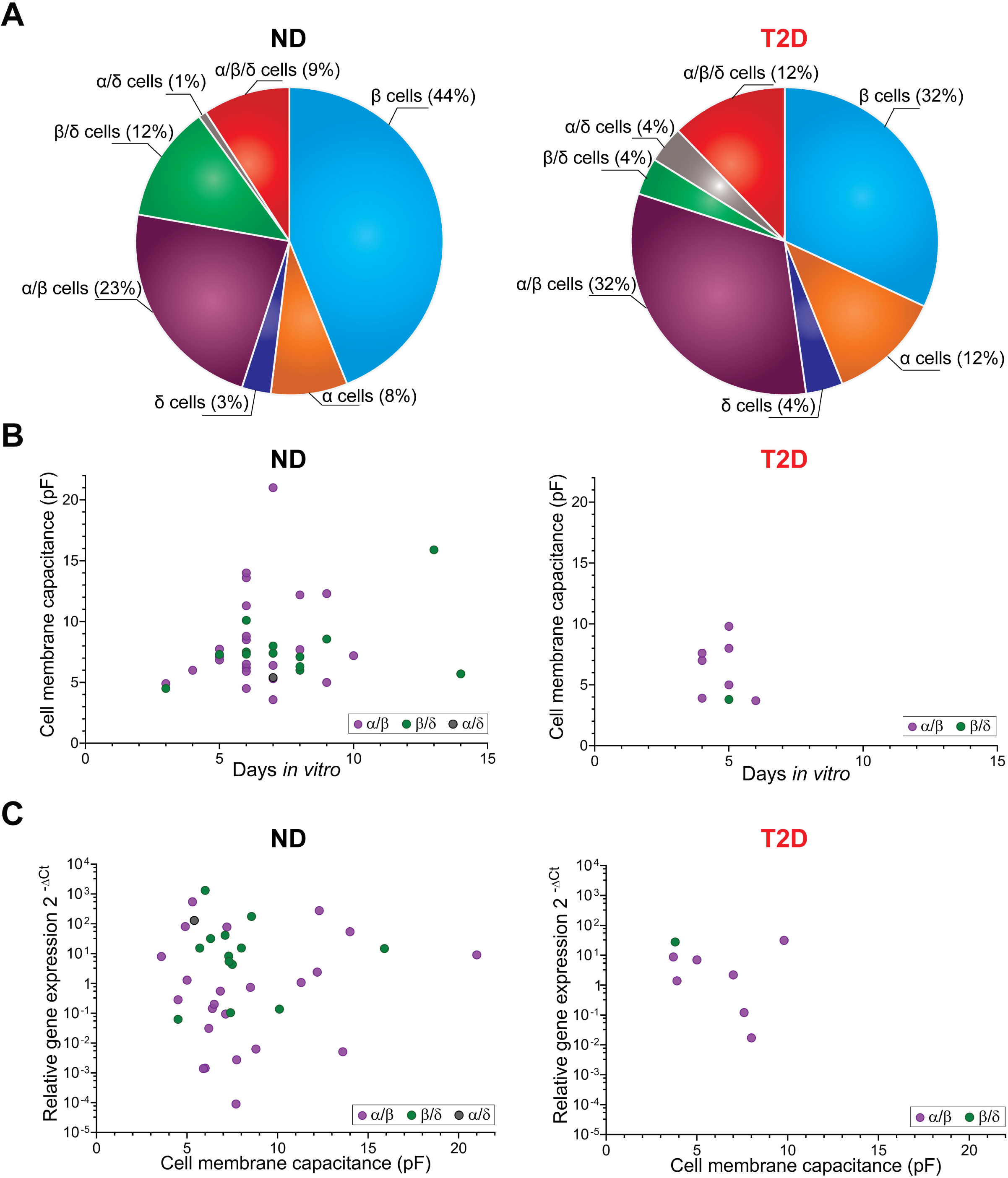
Percentage distribution of single and multiple hormone transcript-expressing cells (**A**) and relations between duration of islet culturing (**B**) and relative gene expression (**C**) versus cell membrane capacitance in intact human pancreatic islets from non-diabetic (ND) and type 2 diabetic (T2D) donors. Relative gene expression in (**C**) is read as the *GCG/INS* expression ratio for mixed-identity α/β cells (magenta circles), *INS/SST* expression ratio for mixed-identity β/δ cells (green circles) and *GCG SST* expression ratio for mixed-identity α/δ cell (gray circle). Correlations neither in (**B**) (Spearman correlation coefficient for ND group r = 0.140, P = 0.410, n = 37; for T2D group r = –0.274, P = 0.514, n = 8), nor in (**C**) (Spearman correlation coefficient for ND group r = –0.019, P = 0.910, n = 37; for T2D group r = –0.238, P = 0.582, n = 8) are revealed.

**Figure 2.**
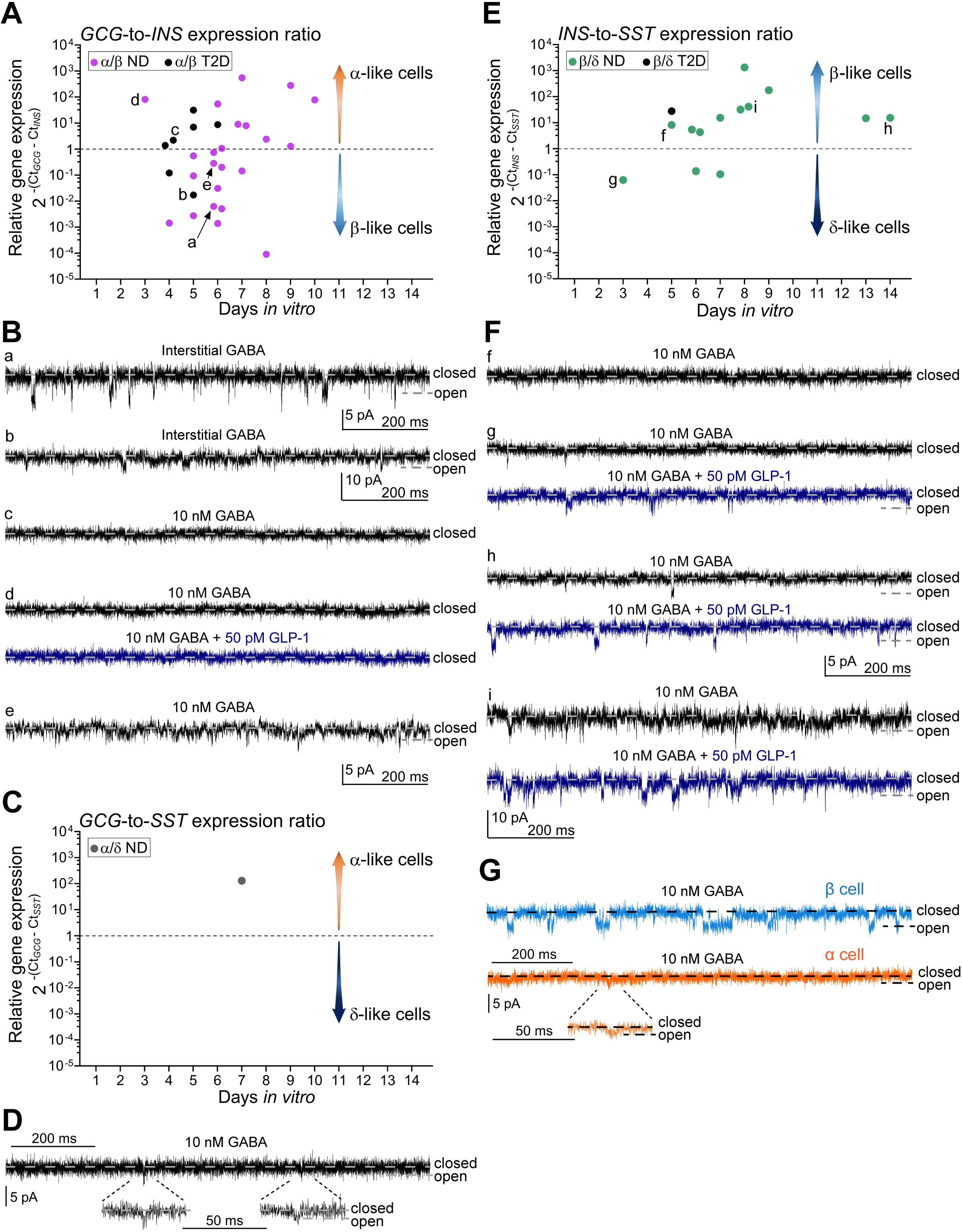
Ratios of hormone mRNA expressions in individual mixed-identity cells with two hormone transcripts and iGABA_A_R-mediated currents in islet cells. (**A**) and (**B**): The scatter dot plot of *GCG/INS* expression ratios in mixed-identity α/β cells (**A**) and representative current recordings through iGABA_A_Rs (**B**) in α/β cells with high (a), low (d) and comparable (c) levels of expression of *INS* relative to the expression level of *GCG*. Dash line at the *GCG/INS* expression ratio = 1 in (**A**) shows equal expression of both hormone transcripts. The higher *GCG/INS* expression ratio, the more α/β cell is α-like (upward arrow); the lower *GCG/INS* expression ratio, the more α/β cell is β-like (downward arrow). (**C**): *GCG SST* expression ratio in an α/δ cell and corresponding recording of iGABA_A_R-mediated current (**D**) in this cell. Two iGABA_A_R single-channel events with low amplitudes are shown at expanded time scale. (**E**): The scatter dot plot of *INS/SST* expression ratios in β/δ cells and representative current recordings through iGABA_A_Rs (**F**) in β/δ cells. For dash line and arrows in (**C**), (**E**) see explanations in (**A**) in context of the respective hormone transcripts. (**G**): Representative recordings showing high activity of single-channel iGABA_A_Rs in a β cell and low activity of single-channel iGABA_A_Rs and lower current amplitudes in an α cell in the presence of 10 nM GABA in ND donors. A single-channel iGABA_A_R opening with low amplitude is shown at expanded time scale. Closed and open states of the single channels are denoted by corresponding dash lines on the recordings (**B**), (**D**), (**F**), (**G**). The scale bars 5 pA and 200 ms are common for the recordings **B**a, c–e and **F**f–h; recordings **B**b and **F**i have vertical scale bar 10 pA. Recordings d, g–i were done in the presence of 10 nM GABA first (black traces), and then 50 pM GLP-1 was added to the extracellular solution in order to examine the potentiation of iGABA_A_Rs via the activation of GLP-1 receptor (blue traces). The recordings in **B**a–b were done without exogenously added GABA, and the rest of recordings were done in the presence of 10 nM GABA.

**Figure 3.**
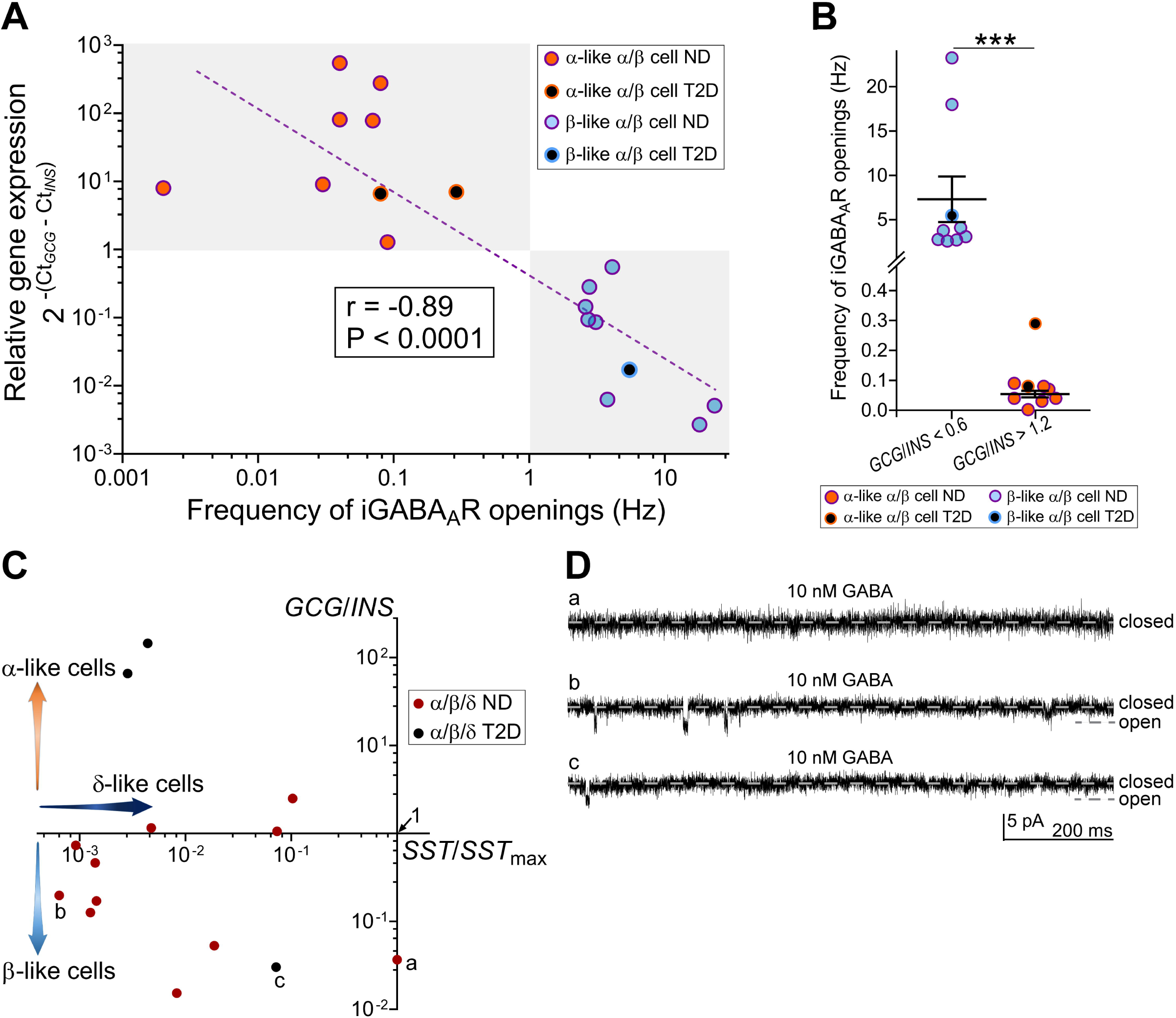
Frequency of single-channel iGABA_A_R openings in mixed-identity α/β cells and combined data for mixed-identity α/β/δ cells. (**A**): Anticorrelation between frequency of single-channel iGABA_A_R openings and the relative *GCG/INS* gene expression in individual mixed-identity α/β cells showing the higher *INS* expression in the α/β cell (= the more mixed-identity cell is a β-like), the higher frequency of the single-channel iGABA_A_R openings in such a cell. Spearman correlation coefficient r = –0.89, P < 0.0001, n = 18. (**B**): The frequency of single-channel iGABA_A_R openings in the α/β cells with the *GCG/INS* ratio between 0.002 and 0.6 (β-like cells, see **A**) is significantly higher than that in α/β cells with the *GCG/INS* ratio between 1.2 and 550 (α-like cells, see **A**). Nonparametric Mann-Whitney test, ***P < 0.0001, n = 9 in the β-like group, n = 8 in the α-like group. The uppermost data point in the α-like group was obtained from T2D donor, detected as outlier by Tukey method and excluded from the comparison. (**C**): Hormone transcript expression in individual mixed-identity α/β/δ cell presented as the *GCG/INS* expression ratio in the cell versus the expression of the *SST* in the same cell divided by the maximal expression of the *SST* among all mixed-identity α/β/δ cells (*SST/SST*_max_). The lower *SST/SST*_max_ ratio, the more negligible *SST* component in mixed-identity α/β/δ cell, and then the cell is considered as mixed-identity α/β cell. Thus, to be e.g. β-like, mixed-identity α/β/δ cell should fall into lower left part of the scatter plot (downward arrow). (**D**): Recordings of iGABA_A_R-mediated currents in three individual mixed-identity α/β/δ cells marked on **C**a–c from non-diabetic donors (a, b) and a type 2 diabetic donor (c). Electrophysiological recordings were done at the V_h_ = –70 mV. Pancreatic islets were exposed to 10 nM GABA.

In rodent islets the cell size normally correlates with the major cell types [15, 16] but the situation is somewhat different for human islet cells where we did not detect any difference in cell size among α, β and δ cells in intact islets [10]. However, it is possible that alterations in size reflect transdifferentiation of one cell-type to another. We, therefore, examined if the mixed-identity cells differed in size or if the time in culture influenced the cells’ diameter. Fig. 1B shows that the different subtypes of cells were similar in size, as determined from cell membrane capacitance measurements, and that the cell size did not correlate with the time in culture after isolation of islets. We further examined if the relative expression level of a pair of hormone transcripts in the mixed-identity cells correlated with the cells size but no correlation was found between these two parameters (Fig. 1C).

### iGABA_A_ receptor-mediated currents in the different subtypes of the mixed-identity cells

We further analysed the recordings of iGABA_A_R-mediated currents in mixed-identity cells in order to examine if a particular subtype of the mixed-identity cells had a characteristic pattern of GABA-activated currents. The single-channel iGABA_A_R currents were recorded in 87% of the mixed-identity cells analysed with both electrophysiological and single-cell RT-PCR techniques. Fig. 2A shows the distribution of the *GCG/INS* expression ratio for individual mixed-identity α/β cells as a function of days in culture after the isolation of pancreas. No effect of time in culture on the *GCG/INS* expression ratio was detected (Spearman correlation coefficient r = 0.22, P = 0.248, n = 30 cells).

Interestingly, recordings from α/β cells with higher relative *INS* expression (corresponding to lower *GCG/INS* values) have higher frequency and larger amplitudes of iGABA_A_R-mediated currents than those with higher *GCG/INS* expression ratio (see Fig. 2Aa, Ba and Ad, Bd). In agreement with this observation, we found strong anticorrelation between relative *GCG/INS* expression levels and single-channel iGABA_A_R opening frequency (Fig. 3A; Spearman correlation coefficient r = –0.89, P < 0.0001, n = 18). Thus, the α-like α/β cells had relative expression levels of 1.2 < *GCG/INS* < 550 and the β-like α/β cells of 0.002 < *GCG/INS* < 0.6 and the difference in frequencies of the single-channel iGABA_A_R openings for α-like α/β cells, 0.054 ± 0.011 Hz, and for β-like α/β cells, 7.30 ± 2.57 Hz, was significantly different (mean ± S.E.M, nonparametric Mann-Whitney test, P < 0.0001, n = 9 in the β-like group, n = 8 in the α-like group; Fig. 3B). This is in line with the patterns of activities of iGABA_A_Rs in single hormone transcript-expressing α and β cells [10] (Fig. 2G) and can be used to discriminate between α- and β-like α/β cells. We also examined if the frequency of single-channel openings of iGABA_A_Rs altered with duration of the islets in culture but no change was detected (Spearman correlation coefficient r = – 0.35, P = 0.15, n = 18).

Glucagon-like peptide-1 (GLP-1) receptors are not expressed in human α cells [3, 17]. Accordingly, in a cell with high *GCG/INS* expression ratio, no potentiation of single-channel iGABA_A_R activity with GLP-1 application was observed (Fig. 2Ad, Bd) consistent with an α cell-like phenotype. Moreover, in a mixed-identity α/δ cell with high expression of *GCG* relative to *SST* (Fig. 2C), we recorded low-frequency single-channel iGABA_A_R-mediated events with low conductance that also corresponds to an α-like cell phenotype (Fig. 2D). In the mixed-identity cells with higher *INS/SST* expression ratios (Fig. 2E), high activity level of the single-channel events with the current amplitudes comparable to those obtained in single-transcript (*INS* only) β cells was generally observed, and the currents were potentiated by GLP-1 application (Fig. 2Fg–i). We also recorded currents through iGABA_A_Rs in mixed-identity α/β/δ cells. The most prominent single-channel iGABA_A_R currents were recorded in cells with the highest *INS* expression among all three hormone transcripts (Fig. 3Cb,c and Db,c). However, the difference in hormone transcripts expression levels varied in the mixed-identity α/β/δ cells and the frequency of the single-channel iGABA_A_R currents was relatively low in these cells (see Fig. 3Db,c and e.g. Fig. 2Ba, Fi).

## Discussion

In recent years, reports have emerged indicating that there are groups of pancreatic islet cells that express more than one hormone transcript [2, 3, 7]. It is possible that these mixed-identity cells have properties different from single hormone transcript-expressing cells. In the current study we analyzed the proportions of single hormone transcript-expressing and mixed-identity cells in islets from non-diabetic and type 2 diabetic donors and further, explored the iGABA_A_R-mediated currents peculiar to a specific mixed-identity cell subtype.

Studies of type 2 diabetes have shown a decrease in the β cell mass and a concomitant augmentation in the number of α cells in islets from type 2 diabetic donors as compared to control subjects [18, 19]. Our cytosome analysis corroborate these results, revealing a decreased probability of identifying single-hormone *INS*-expressing β cells in islets from type 2 diabetic donors compared to islets from non-diabetic subjects. In contrast, the probability of identifying cells containing the *GCG* increased and, in particular, the percentage of single-hormone *GCG*-expressing α cells increased in islets from type 2 diabetic donors. Whether this change is a cause or a consequence of the disease remains to be determined. Interestingly, different subtypes of the mixed-identity cells express hormone transcripts at variable levels, and several combinations exist. Importantly, however, no systematic change in the *GCG/INS* expression level was observed for the cells during the 10 days after isolation from the donors.

Apparently, the mixed-identity cells have distinct intracellular regulatory mechanisms governing particular hormone transcript expression. These cells may also differ in iGABA_A_R subunit composition and their expression levels that will be reflected in different patterns of single-channel iGABA_A_R openings.

We have previously characterized the functional properties of iGABA_A_R in human α and β cells [10]. Here, in mixed-identity α/β cells, we found that cells having higher *GCG/INS* expression ratio correlated with no or low single-channel iGABA_A_R opening frequency and low-amplitude single-channel events and no response to GLP-1 application. This pattern of activity is very much similar to the behavior of iGABA_A_Rs in single-hormone *GCG*-expressing α cells [10]. On the other hand, mixed-identity α/β cells with lower *GCG/INS* expression ratio had activity similar to single-hormone *INS*-expressing β cells [10] with higher frequency and larger amplitudes of single-channel iGABA_A_R openings. In the majority of the mixed-identity β/δ cells we found that the *INS* expression level was higher than that for somatostatin and the pattern of activity of single-channel iGABA_A_Rs was similar to the activity pattern in β cells. Together, the results identify the iGABA_A_Rs as a functional marker of the physiological identity of the mixed-identity cell subtype.

The explanations for the existence of mixed-identity cells in the human pancreatic islets may be many. The human islet is a plastic structure [1, 20, 21] and numerous factors [4, 5, 22], including GABA [23, 24] may influence the signatures of the cells. The cell-type determination has been proposed to take place during development [25] or alter due to dedifferentiation [22] or intentional reprogramming [7]. Further studies are required to identify factors and conditions regulating the cell-type identity [26].

The GABA signaling system is an integral part of the normal human pancreatic islet physiology [8, 27]. If pancreatic islet GABA concentration changes out of the physiological range, it may impair proper insulin and glucagon secretion, potentially alter cell fate [23, 24, 28] and eventually contribute to pathogenesis of type 2 diabetes. Moreover, interstitial GABA has also been proposed to inhibit cytotoxic immune cells entering the islets and is of potential importance for both type 1 and type 2 diabetes [28-30].

In conclusion, our results show that iGABA_A_R activity predicts the phenotype of the mixed-identity cells. Better understanding of the GABA signalling system effects in the human pancreatic islets will be valuable and may assist in unravelling the relationship between the α and the β cells plus, potentially, how the intrinsic potential for regeneration of the β cell mass comes about.

## Acknowledgements

The authors thank the Nordic Network for Clinical Islet Transplantation for generous providing human pancreatic islets. This work was supported by Swedish Research Council grants (grant numbers 521-2009-4021, 521-2012-1789, 2015-02417 to B.B.), Diabetes Wellness, Swedish Diabetes Foundation, the Novo Nordisk Foundation, The Swedish Children’s Diabetes Foundation, Family Ernfors Foundation, The strategic grant consortium Excellence of Diabetes Research in Sweden (EXODIAB). S.V.K. was supported by E. Wessler’s foundation and Astrid Karlsson’s foundation for medical research (Uppsala University) as well as Thurings Foundation.

## Author Contributions

S.V.K., Z.J. and B.B. designed experiments; S.V.K. and Z.J. performed experiments; S.V.K. and Z.J. analysed data; S.V.K. made the figures; S.V.K. and B.B. wrote the manuscript. B.B. is the guarantor of this work and, as such, had full access to all the data in the study and takes responsibility for the integrity of the data and the accuracy of the data analysis.

## Additional Information

### Competing Interests

B.B. has filed two patent applications based on GABA and GABA_A_ receptors function. S.V.K. and Z.J. have no conflict of interests to disclose.

